# Non-canonical IL-2 signaling facilitates peripheral regulatory T cell differentiation

**DOI:** 10.64898/2026.10.07.756301

**Authors:** Amelia Meecham, Sarah Hyun Ji Kim, Mikhael Haruo Fernandes de Lima, Michelle Tang, Bryan Liao, Samuel Chan, Ho-sup Lee, Li-Fan Lu, Ziyue Zhang, Caroline Jefferies, Monica Guma, Miguel-Alejandro Lopez-Ramirez, Mark H. Ginsberg

## Abstract

Non-canonical IL-2 signaling, termed **M**onoclonal antibody or **M**atrix-directed **A**lternative **C**ytokine **S**ignaling (MACS), is mediated by a CD25–chemokine receptor complex and operates in concert with the canonical JAK-STAT5 pathway; however, its role in T cell homeostasis remains poorly defined. Here, we uncover a role for MACS in regulatory T cell (Treg) differentiation and functional specification. We generated *Il2*^E66K*/E66K*^ mice in which CD25–CCR7 assembly is disrupted and IL-2-induced STAT5 phosphorylation is preserved. Immunophenotyping analysis by flow cytometry revealed that MACS deficiency reduces the accumulation of Nrp1^-^ Foxp3^+^ Tregs in gut-associated lymphoid tissues from 24-weeks-old *Il2*^E66K/E66K^ mutant mice. Single-cell transcriptomics, functional suppression assays, and *in vitro* differentiation revealed that MACS engagement drives Treg differentiation and sustains effector Treg programs—marked by robust IL-10 expression—in part via an AP-1-dependent transcriptional network. These findings establish IL-2 MACS as a distinct signal directing peripheral immune tolerance and highlight a potential node for targeted immunomodulation.

**One Sentence Summary:** Non-canonical IL-2 signaling facilitates gut regulatory T cell development and suppressive function.

## INTRODUCTION

Interleukin-2 (IL-2) has an essential role in the development, maintenance, and function of regulatory T cells (Tregs) (1, 2). Tregs are a subset of CD4⁺ T cells characterized by their expression of the transcription factor Foxp3 and high levels of CD25 (IL-2Rα)(3). They are indispensable for maintaining immune homeostasis and preventing autoimmunity. Treg differentiation, proliferation, and suppressive capacity are dependent on IL-2 availability and signaling (4, 5). Canonical IL-2 signaling is initiated by the formation of a high-affinity IL-2 receptor complex, comprising IL-2Rα (CD25), IL-2Rβ (CD122), and the common γ-chain (CD132)(6). IL-2 binds to CD25, enabling recruitment of the β and γ subunits to form a high affinity trimeric receptor that activates Janus kinases JAK1 and JAK3, leading to phosphorylation of STAT5 and subsequent transcriptional activation of IL-2–responsive genes(7, 8).

We previously reported that IL-2 can also signal through a complementary pathway termed MACS (**M**onoclonal Antibody or **M**atrix-directed **A**lternative **C**ytokine Signaling)(9). This pathway is initiated when CD25-IL-2 forms a complex with certain chemokine receptors(10), including CCR7, leading to Gα_i/o_-dependent integrin activation. IL-2 MACS can be engaged by heparan sulfate (HS) or the anti-CD25 antibody, PC61, leading to enhanced integrin activation and improved Treg suppressive function(9). Moreover, administration of a recombinant effector-incompetent PC61, designated mPC61, increased Treg abundance in the spinal cord and improved disease outcome in murine experimental autoimmune encephalomyelitis. Integrins mediate CD4⁺ T cell function, trafficking and expansion, including in Tregs (11–13). In Tregs, high expression of integrins such as α4β1, αLβ2 and α4β7 facilitates migration to lymph tissue and peripheral sites of inflammation (14, 15). Integrins stabilize interactions between T cells and the microenvironment by binding to ligands in the extracellular matrix and on the surface of antigen presenting cells (APCs)(16), promoting tissue retention and immune function(14).

Structure-based design combined with a medium throughput screening strategy identified a point mutation in human IL-2 that disrupts the capacity of IL-2 to drive assembly of a complex of CD25 with CCR7 or CXCR4 and resulting integrin activation (10). Importantly, this mutation preserved IL-2’s capacity to activate canonical IL-2 signaling as assessed by STAT 5 phosphorylation. Here we have extended this work by identifying and characterizing a mouse IL-2 point mutant (lL-2(E66K)) with similar functional effects and used that information to create mice with reduced IL-2 MACS. We find that these mice exhibit a reduction in the aging-associated increase in Treg abundance in the gut-associated lymphoid tissue (GALT). Single cell transcriptomic analysis of mesenteric lymph nodes from these mice revealed reduced expression of the suppressive cytokine, IL-10 mRNA associated with reduced IL-10 abundance in Tregs and corresponding decreased suppressive function. Furthermore, enforced IL-2 MACS increased in vitr*o* Treg differentiation and was impaired in *Il2^E66K/E66K^* CD4^+^ cells. In sum, these data reveal that MACS contributes to IL-2’s capacity to enable the differentiation and function of pTregs that maintain peripheral tolerance.

## RESULTS

### Murine Il-2 (E66K) disrupts MACS while preserving canonical signaling

The human IL-2(E52K) mutation abrogates antibody-induced assembly of the CD25–IL-2–CCR7 complex and associated MACS pathway dependent integrin activation(10). We therefore sought to determine whether an analogous substitution in murine IL-2 phenocopies the effects of the human IL-2(E52K) mutation. The alignment of human (hIL-2) and mouse IL-2 (mIL-2) protein sequences (Figure 1A) led to the design and construction of mIL-2(E66K). Like hIL-2(E52K), mIL-2(E66K) failed to support antibody and HS-induced association of mCD25 with CCR7 (Fig. 1B).

**Figure 1.**
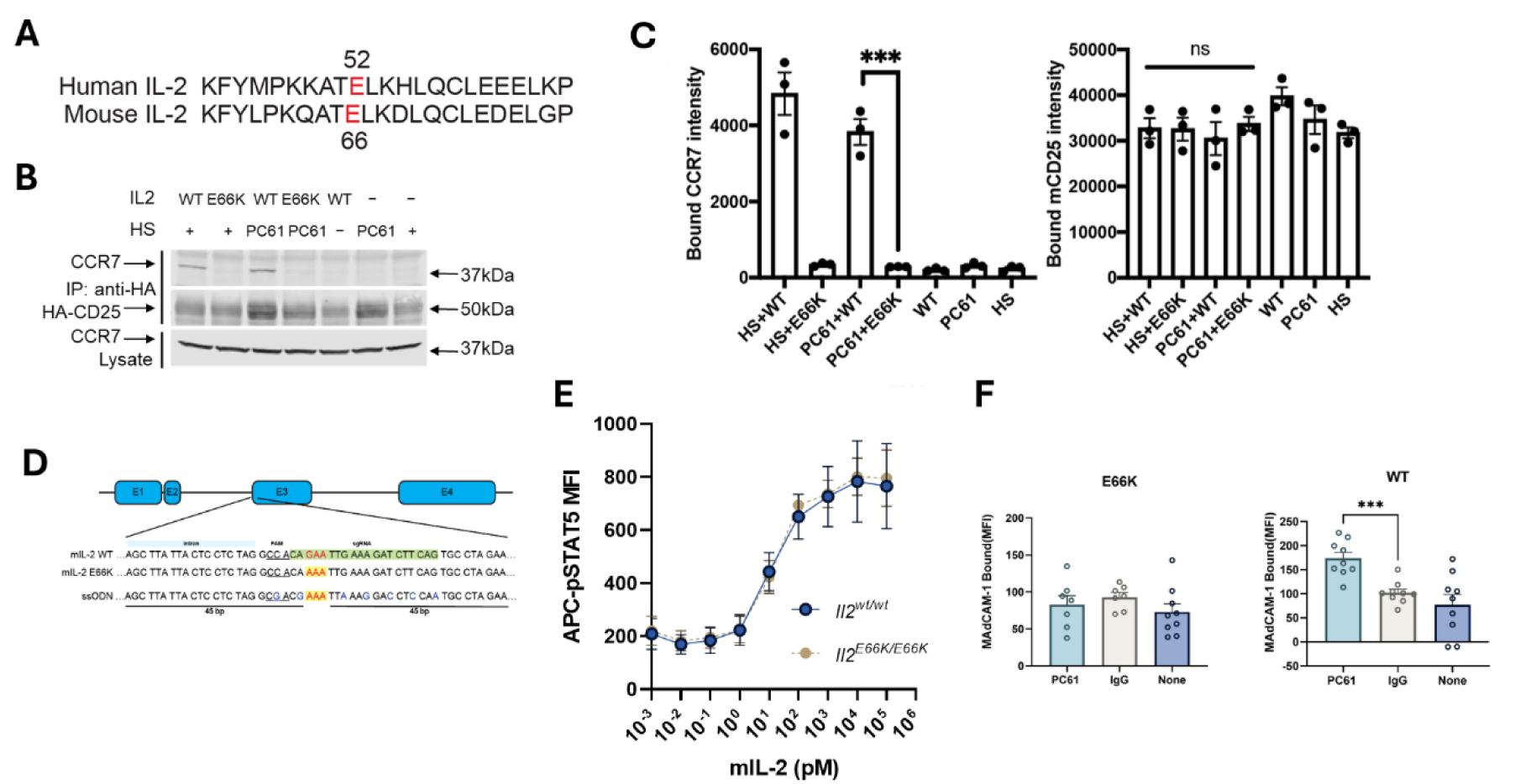
mIL-2(E66K) disrupts CD25–CCR7 association and preserves canonical IL-2 signaling. **(A)** mIL-2(E66K) was identified based on an alignment with hIL-2 wherein the hIL-2(E52K) mutations abolishes MACS via CCR7 and CXCR4. **(B&C)** mIL-2(E66K) blocks CD25-CCR7 association. HEK293T HA-mCD25 cells or for HS-dependent complex formation, lysed membrane fractions of HA-mCD25 HEK293T cells were incubated with 5 µg/mL PC61 in the presence or absence (-) of 1 µg/mL IL-2 (WT) or IL-2(E66K) (E66K) and after lysis, mHA-CD25 was immunoprecipitated and fractionated by SDS-PAGE. Immunoblots (B) were probed with either anti-HA or anti-CCR7 for captured mCD25 or associated CCR7 and results from 3 independent experiments were quantified in (C). *p<0.05, **: p<0.01, ***: p<0.001, ****: p<0.0001 by one-way ANOVA. **(D)** Schematic of the ssODN repair template used for CRISPR–Cas9 editing of the mouse *Il2* locus to introduce the E66K mutation **(E)** Foxp3-GFP^+^ Tregs from WT or *Il2*^E66K/E66K^ mice were incubated with increasing concentrations of IL-2(WT) or IL-2(E66K), respectively, for 30 min at 37°C followed analysis of phospho-STAT5 staining by flow cytometry. **(F)** CD4^+^ T cells from Foxp3-GFP WT or *Il2*^E66K/E66K^ were incubated with 5 μg/mL MAdCAM-1 Fc for 30 min at 37°C with 0.5 μg/mL IL-2 or IL-2 (E66K) respectively in the presence of 10 μg/mL PC61 or control IgG. In parallel 10 mM EDTA was added in each condition to measure non-specific binding. After washing cells were stained with APC-anti-human IgG Fc and geometric mean fluorescence intensity (MFI) was determined on Foxp3-GFP^+^ Tregs. Specific MAdCAM-1 binding (ΔEDTA mean fluorescence intensity, MFI) was measured as a readout of integrin activation. Statistical significance was determined using two-way repeated measures ANOVA with genotype and treatment as factors (genotype × treatment interaction p = 0.0041), followed by post hoc comparisons within each genotype as indicated. ***p < 0.001; ns, not significant.

To investigate the physiological role of the MACS pathway, we created a *Il2*^E66K/E66K^ transgenic mouse using CRISPR–Cas9–mediated genome editing. A single-stranded oligodeoxynucleotide (ssODN) repair template was designed to create the desired codon change (GAA→AAA, Glu66→Lys) flanked by wild-type mIL-2 sequence homology arms (Figure 1C). The ssODN also included nearby silent synonymous nucleotide substitutions to disrupt the Cas9 guide/PAM site and provide a genotyping marker.

Critically, IL-2(E66K) did not compromise the canonical IL-2 signaling pathway in murine Tregs as assessed by STAT5 phosphorylation (Figure 1D). MAdCAM-1 binding was used to assess downstream MACS in WT and *Il2*^E66K/E66K^ Tregs following stimulation with PC61. In WT Treg, PC61 significantly increased MAdCAM-1 binding, whereas no such increase was observed in *Il2*^E66K/E66K^ cells (Fig. 1E). Two-way repeated measures ANOVA revealed a significant genotype × treatment interaction (p = 0.0041), indicating that PC61-induced integrin activation is dependent on intact IL-2 MACS signaling. These data establish IL-2(E66K) as a mouse model for analyzing MACS dependent functions of IL-2.

### Reduced Tregs in the GALT of aged *Il2^E66K/E66^*^K^ mice

*Il2*^E66K/E66K^ mice were fertile, viable, and showed no overt signs of autoimmune disease. Complete blood counts of whole blood showed no significant differences between *Il2*^E66K/E66K^ and its WT littermate controls (Supplementary Figure 1). Spleens, secondary lymph nodes (LN), Peyer’s patches and colon from *Il2*^E66K/E66K^ and *Il2*^WT/WT^ mice were collected to immunophenotype CD4⁺ T cells and Tregs. Aged *Il2*^E66K/E66K^ mice (6 months or older) showed a significant reduction in the percentage of Foxp3^+^ Tregs present in the gut associated lymphoid tissue (GALT), specifically, mesenteric LN (mLN) and lamina propria lymphocytes from colon (cLPL) (Figure 2B) whereas, no such reduction was observed in younger (e.g. 12 week old) mice.

**Figure 2.**
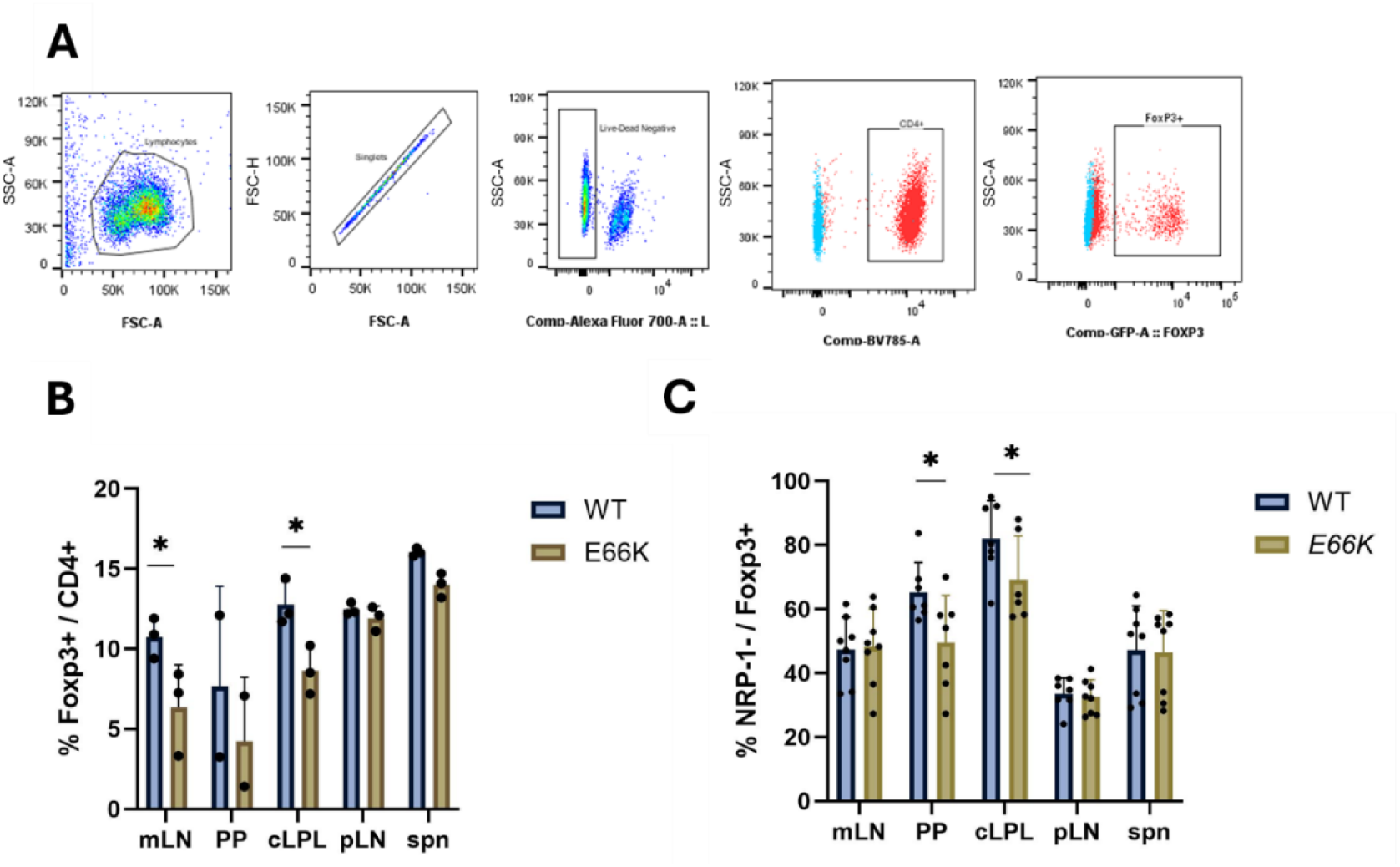
Reduced Treg in the GALT of aged *Il2*^E66K/E66K^ mice. **(A)** Representative flow cytometry plots showing sequential gating of mesenteric lymph node (LN) cells. Lymphocytes were first identified based on forward and side scatter, followed by exclusion of doublets (singlets) and dead cells (Live/Dead negative). CD4⁺ T cells were then gated, and Foxp3-GFP⁺ cells were identified within the CD4⁺ population. Blue points indicate fluorescence minus one (FMO) controls used to define gating boundaries for CD4 and Foxp3. Gates were applied consistently across all samples. **(B)** Immunophenotyping of CD4⁺ T cells from secondary LN of aged *Il2*^E66K/E66K^ mice reveals reduction in % Tregs in GALT. spn: spleen; pLN: peripheral LN; mLN: mesenteric LN; PP: Peyer’s patches; cLPL: colon lamina propria lymphocytes. (n=3) **(C)** Immunophenotyping of CD4⁺ T cells from the same peripheral tissues in >6-month-old *Il2*^E66K/E66K^ mice reveals a selective reduction in Nrp1⁻ Tregs compared with littermate controls. All statistics are 2-way ANOVA with Tukey correction, mean ± SD. ns: not significant; *: p<0.05

There are two main origins of Tregs, thymic and peripheral Tregs. Thymic Tregs (tTregs) are originally differentiated and expanded in the thymus, mediated by the specificity of T cell antigen receptors (TCRs) to self-antigens(*17*). In contrast, peripheral Tregs (pTregs) are differentiated from naïve CD4⁺ T cells extra-thymically(*18*). To partially distinguish between tTregs and pTregs, we analyzed Nrp-1 expression in GALT Tregs(*19*). Aged *Il2*^E66K/E66K^ mice (>6 months) exhibited significantly reduced frequencies of Nrp-1⁻ Tregs in Peyer’s patches and colonic lamina propria (Fig. 2C), indicating a defect in sustaining the pool of Nrp1⁻ Foxp3^+^ Tregs.

### Single-cell transcriptomics identifies compromised effector Treg function in Il2^E66K/E66K^ mice

We performed single-cell RNA sequencing (scRNA-seq) on sorted CD4⁺ T cells from the mLNs of aged *Il2*^E66K/E66K^ or *Il2*^WT/WT^ 24 week old mice (summary statistics and quality control metrics can be found in Supplementary Fig. 2). Unsupervised clustering and UMAP visualization identified major CD4⁺ T cell subsets (Figure 3A), which were annotated using established marker genes (Supplementary Figure 3) as described (*20*). The largest cluster in both genotypes corresponded to naïve cells, with a modest decrease in frequency in E66K mice (52.30% WT vs. 46.94% E66K). Although overall Treg abundance was comparable between groups, total CD4^+^ T cells from *Il2*^E66K/E66K^ mice exhibited reduced expression of *Foxp3, Il2ra (CD25)* and *Ctla4*, indicating compromised Treg identity despite preserved abundance (Figure 3B). In total, 138 genes were differentially expressed (p adjusted value <0.05 and logFc >1.5 or <-1.5), 133 genes were downregulated in *Il2*^E66K/E66K^ cells, and the remaining 5 were upregulated. A summary of the top differentially expressed genes (DEGs) can be found in Figure 3C.

**Figure 3.**
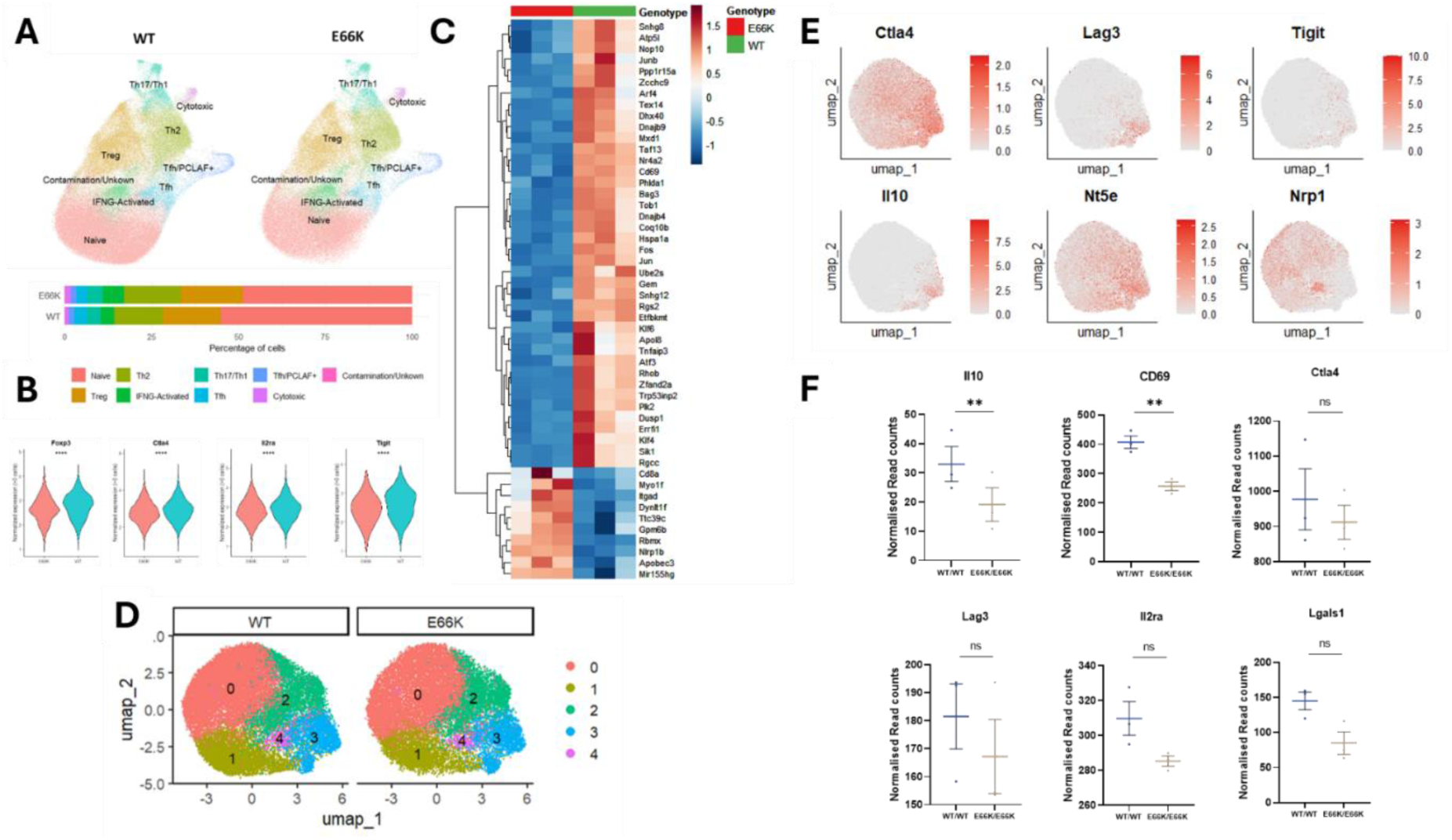
Single-cell RNA sequencing of mesenteric lymph node CD4⁺ T cells reveal reduced Il-10 expression in an effector Treg subset in *Il2*^E66K/E66K^ mice. **(A)** UMAP visualization of scRNA-seq profiles from mesenteric lymph node CD4⁺ T cells from wild-type and *Il2*^E66K/E66K^ mice (n=3 per genotype), with major cell types annotated; accompanying bar plot shows the relative distribution of cell types by genotype**. (B)** Violin plots showing normalized single-cell expression of key Treg-associated genes (*Foxp3, Ctla4, Il2ra, and Tigit*) across all CD4⁺ T cells from mesenteric lymph nodes of WT and *Il2*^E66K/E66K^ mice. Statistical significance was assessed using a Wilcoxon rank-sum test comparing all single cells between genotypes. **(C)** Heat map of pseudobulk differential expression across all identified CD4⁺ T cell populations, highlighting the top differentially expressed genes (DEGs) between WT and *Il2*^E66K/E66K^ mice. **(D)** Subclustering of the Treg compartment visualized by UMAP and split by genotype. (E) Feature plots of Treg-associated transcripts (*Ctla4, Tigit, Lag3, Il10, Nt5e, and Nrp1*) identify a suppressive effector Treg subcluster (subcluster 3**) (F)** Pseudobulk differential expression analysis within subcluster 3 demonstrates reduced Il10 expression in *Il2*^E66K/E66K^ compared with WT samples. Statistical significance is indicated as follows: *P < 0.05; **P < 0.01; ***P < 0.001; ****P < 0.0001.

We noted that the reduction in Treg frequency observed by flow cytometry was not recapitulated in the scRNA-seq analysis. This likely reflects differences in how Tregs are annotated across these platforms. In flow cytometry, Tregs are identified using a binary threshold of Foxp3 expression, classifying cells as Foxp3⁺ or Foxp3⁻. In contrast, scRNA-seq defines Tregs based on transcriptional clustering and gene expression signatures, which may include cells with lower or heterogeneous *Foxp3* expression(*21*). Consistent with this, examination of *Foxp3* expression across all CD4⁺ T cells revealed a reduction in median expression in *Il2*^E66K/E66K^ cells, even in the absence of a clear shift in cluster-defined Treg frequency (Fig. 3B).

To gain further insight, we focused on the Treg-annotated cluster and performed subclustering (Figure 3D). This analysis resolved the Treg compartment into five distinct clusters, one of which (Cluster 3) was characterized by elevated expression of *Tigit, Ctla4, Lag3, Il10* and *Nt5e*, together with low expression of *Nrp1* and *Helios*, consistent with a peripherally differentiated effector Treg (eTreg) phenotype (Figure 3E) (*19, 21–24*). Pseudobulk differential expression analysis of this eTreg subset revealed that *Il2*^E66K/E66K^ mice exhibited reduced expression of *Il10* compared to wild-type controls, suggesting impaired eTreg suppressive function (Figure 3F). Notably, in our previous work (*9*), activation of the MACS pathway using PC61 antibody led to increased IL-10 expression and enhanced Treg suppressive capacity, supporting the idea that this signaling axis promotes eTreg function.

To extend these findings to protein expression, we performed flow cytometric analyses of mesenteric lymph node CD4⁺ T cells. Total CD4⁺ T cells were isolated directly from *Il2*^E66K/E66K^ and *Il2*^WT/WT^ mice, confirming a reduced frequency of Foxp3⁺ Tregs within the CD4⁺ compartment in *Il2*^E66K/E66K^ samples. Intracellular cytokine staining demonstrated significantly lower IL-10 expression in *Il2*^E66K/E66K^ Tregs compared with WT controls (Fig. 4C). Consistent with a potential impairment of suppressive programs, expression of additional inhibitory molecules, including TIGIT and CTLA-4, also trended lower in *Il2*^E66K/E66K^ Tregs.

**Figure 4.**
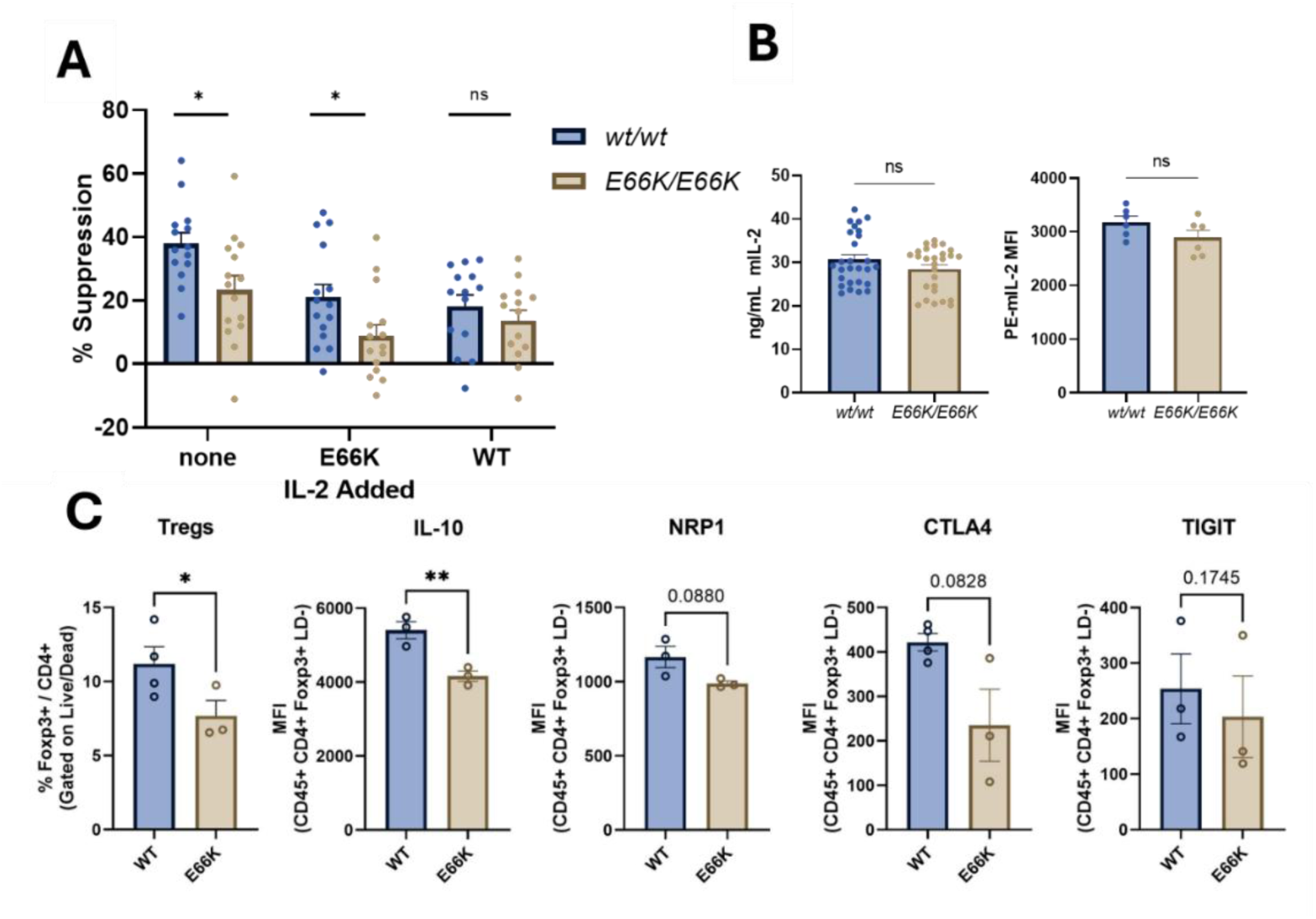
*Il2*^E66K^ Tregs exhibit impaired suppressive function and reduced Il10 expression. **(A)** Suppressive function of WT (blue) and *Il2*^E66K/E66K^ (gold) Foxp3-GFP⁺ Tregs assessed in a CD4⁺ T cell proliferation assay. FACS-sorted naïve CD4⁺CD62L^hi^ FOXP3-GFP⁻ responder T cells were labeled with eFluor670 and cultured (5 × 10⁴ cells/well) with anti-CD3 (5 µg/mL) and BMDCs (3 × 10³ cells/well), in the absence or presence of FACS-sorted Tregs (6.25 × 10³ cells/well; responder:Treg ratio 8:1. Where indicated, exogenous WT or IL-2(E66K) (6ng/ml) was added. After 4 days, responder cell proliferation was quantified by eFluor670 dilution by flow cytometry, and division index was calculated. Percent suppression was determined relative to cultures lacking Tregs. Each point represents the mean of triplicate wells from one experiment. **(B)** IL-2 production by CD4⁺ T cells following *in vitro* stimulation, quantified by ELISA (left) and intracellular cytokine staining (right). Data show IL-2 concentration in culture supernatants and mean fluorescence intensity (MFI) of intracellular IL-2. **(C)** Flow-cytometric analysis of mesenteric lymph nodes from WT and *Il2*^E66K/E66K^ >24 week old mice showing total Treg frequency among CD4⁺ cells, and expression of IL-10, TIGIT, and CTLA-4, measured as MFI within the Foxp3-GFP⁺ gate. Each dot represents an individual mouse. Bars indicate mean ± SE Statistical significance was assessed using two-tailed tests as indicated; *p < 0.05, *p < 0.01; ns, not significant.

We next assessed whether the reduced expression of IL-10 in *Il2*^E66K/E66K^ eTregs translated to impaired suppressive function. In vitro suppression assays were performed by co-culturing eFluor670-labeled genotype-matched responder T cells with anti-CD3 and bone marrow-derived dendritic cells (BMDCs) in the presence of Tregs isolated from *Il2*^E66K/E66K^ or *Il2*^WT/WT^ mice. Consistent with our previous findings of reduced IL-10 expression, IL-2(E66K) Tregs exhibited significantly reduced suppression of responder cell proliferation (Figure 4A). Furthermore, to test whether exogenous IL-2 could correct the defect in IL-2(E66K) Tregs, we added either IL-2 or IL-2(E66K) to the culture. Supplementation with WT IL-2 reduced overall suppression and nearly eliminated the difference between WT and E66K Tregs, suggesting that high levels of wild type IL-2 mask the intrinsic defect. In contrast, addition of IL-2(E66K), although also reducing suppression, maintained the relative functional impairment of IL-2(E66K) Tregs. Importantly, IL-2 production and secretion by *Il2*^E66K/E66K^ or *Il2*^WT/WT^ CD4⁺ T cells were quantified by intracellular staining and ELISA of culture supernatants and were not significantly different (Figure 4B). Thus, impairment of IL-2 MACS in the *Il2^E66K/E66K^*mouse is associated with age related reduction in GALT Treg abundance, reduced expression of suppressive genes, and reduced suppressor function.

### Enforcing MACS enhances iTreg generation from wild type but not *Il2^E66K/E66K^* CD4^+^ cells

The reduction of Nrp1⁻ Tregs in the gut of older *Il2^E66K/E66K^*mice resembles the pTreg development defect of *Foxp3^ΔCNS1^* mice described by Rudensky and colleagues(*25, 26*). Mice lacking CNS1 develop normally and maintain intact thymic Tregs, yet exhibit a marked reduction in pTregs, particularly within GALT. Despite this defect, Foxp3^ΔCNS1^ mice do not display systemic autoimmunity, indicating that CNS1 is dispensable for tTreg development but important for pTreg generation. The similarity of phenotype between *Il2^E66K/E66K^*and *Foxp3^ΔCNS1^* mice raises the possibility that the MACS pathway signals via the CNS1-dependent promoter region of Foxp3 and drives pTreg differentiation.

To test whether activation of the MACS pathway directly affects Treg differentiation, naïve CD4⁺ T cells were isolated from *Il2^WT/WT^Foxp3^GFP^* and *Il2^E66K/E66K^Foxp3^GFP^* splenocytes and cultured for 4 days to induce Treg differentiation using integrin αvβ8-coated plates (*27*). Enforcement of MACS signaling by addition of mPC61 (*9*) led to a ∼2 fold increase in Foxp3^+^ Tregs in comparison to isotype treated control cultures (Fig. 5A) whereas no such increase was seen with , *Il2^E66K/E66K^* CD4^+^ naïve T cells. There was thus a significant difference observed in the capacity of mPC61 to induce iTreg differentiation between the *Il2^wt/wt^* and *Il2^E66K/E66K^* cultures (% increase in Foxp3-GFP expressing cells) (Fig. 5B).

**Figure 5.**
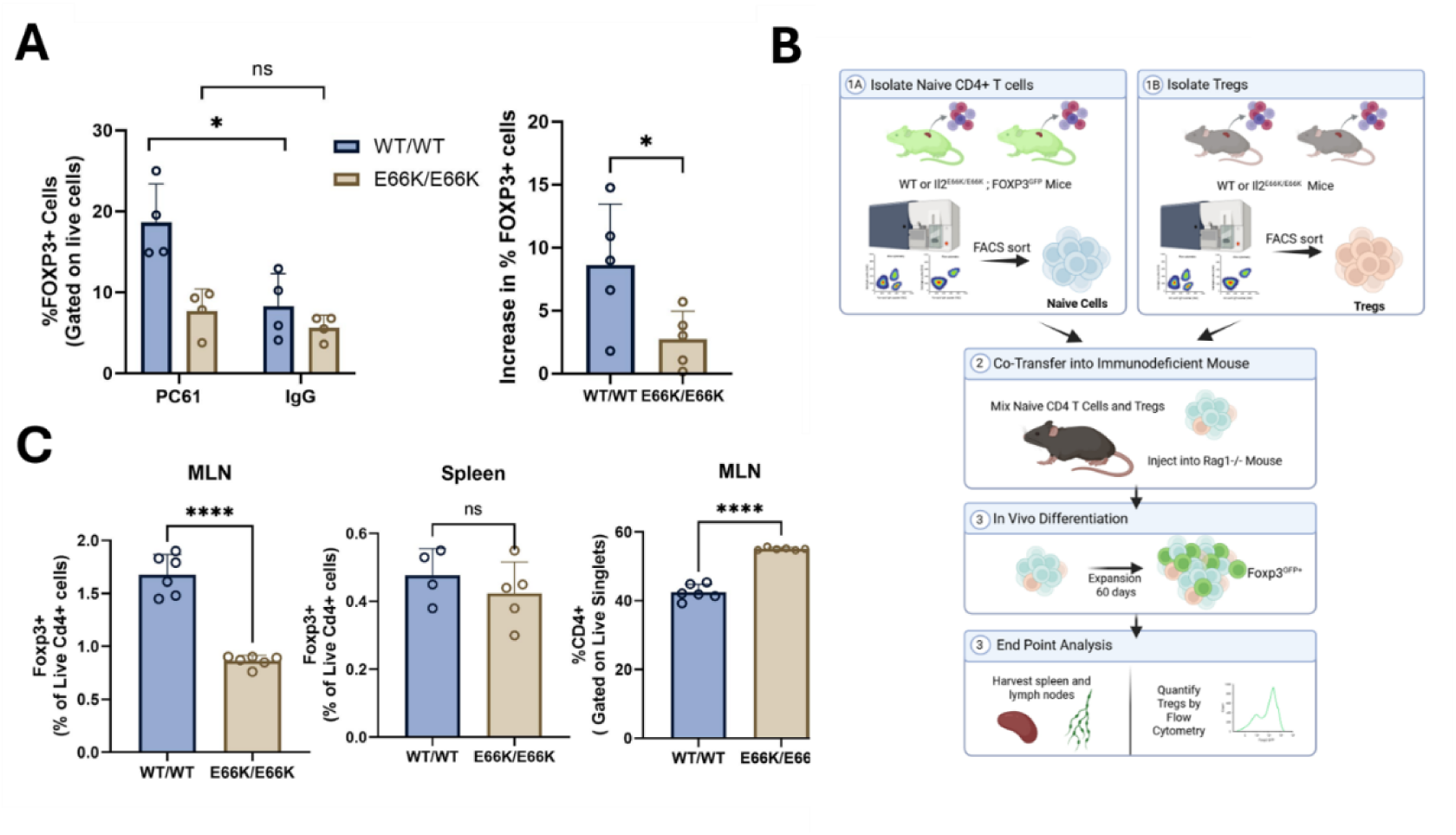
Disruption of MACS signaling impairs peripheral Treg differentiation from naïve CD4⁺ T cells. **(A)** Frequency of Foxp3-GFP⁺ iTregs generated from WT and *Il2*^E66K/E66K^ naïve CD4⁺ T cells in the presence of mPC61 or isotype control, determined by flow cytometry. **(B)** Schematic of *Rag1⁻/⁻* adoptive transfer experiment. Naïve CD4⁺ T cells (CD4⁺ Foxp3-GFP⁻) were isolated from WT or *Il2*^E66K/E66K^ Foxp3-GFP mice in parallel with CD4⁺ Foxp3⁺ Tregs purified from genotype-matched Foxp3-GFP^-^donors. Cells were mixed at defined ratios and adoptively transferred into Rag1⁻/⁻ recipient mice. Following homeostatic expansion, naïve CD4⁺ T cells differentiated into Foxp3-GFP⁺ peripheral Tregs (pTregs) in vivo. Mesenteric lymph nodes and spleen were harvested and analyzed by flow cytometry to quantify CD4⁺ T cells and Foxp3-GFP⁺ cells within the CD4⁺ compartment. **(C)** Frequency of CD4⁺ T cells among total lymphocytes (left) and frequency of Foxp3-GFP⁺ cells within the CD4⁺ compartment in mesenteric lymph nodes and spleen following adoptive transfer. *Il2*^E66K/E66K^ CD4⁺ T cells exhibit reduced Foxp3-GFP⁺ Treg abundance compared to WT controls. Three technical reads were averaged per mouse, and group differences were assessed using an unpaired two-tailed t test, with n = 2 mice per group (p<0.05).

To test further the hypothesis that MACS regulates pTreg differentiation, we performed an adoptive transfer experiment using *Rag1^-/-^* recipient mice. Naïve CD4^+^ cells from *Il2^WT/WT^Foxp3^GFP^* and *Il2^E66K/E66K^Foxp3^GFP^* mice were transferred into *Rag1^-/-^* recipients that were sacrificed after 60 days (Figure 5B). Consistent with our *in vitro* findings, *Il2^WT/WT^Foxp3^GFP^* donor cells led to greater GALT Treg abundance *in vivo* compared to *Il2^E66K/E66K^Foxp3^GFP^* cells, as measured by frequency of Foxp3-GFP^+^ cells in mesenteric lymph nodes (Figure 5C). In contrast, no significant difference was observed in Foxp3-GFP^+^ cells isolated from the spleen.

### MACS-dependent increase in iTreg differentiation occurs without increased TGF-β signaling

TGF-β signaling leads to phosphorylation of Smad3 (*28*) resulting to its binding to the *Foxp3* CNS1 enhancer that drives Treg differentiation. To determine whether increased TGF-β signaling accounts for the MACS-dependent increase in pTreg differentiation we assessed PC61 enhanced Smad2/3 phosphorylation following mPC61 treatment. Naïve CD4⁺ T cells from *Il2*^WT/WT^ mice were cultured under iTreg differentiation conditions in the presence of either mPC61 or isotype control antibody and whole-cell lysates were collected at 48, 72 and 96 hours for Western blot analysis (Figure 6A). Densitometric quantification revealed no increase in phospho-Smad2/3 relative to total Smad2/3 levels in mPC61-treated samples compared to control. In addition, we compared TGF-β signaling between *Il2*^WT/WT^ and *Il2*^E66K/E66K^ T cells under these conditions, with no differences observed in phospho-Smad2/3 abundance (Fig. 6A). Furthermore, bulk RNA sequencing was performed on CD4⁺ T cells cultured for 4 days under iTreg-inducing conditions with mPC61 or IgG control. As expected, cells cultured under Treg differentiation conditions displayed robust transcriptional activation of the TGF-β signaling axis compared to control (Figure 6B). However, differential expression analysis revealed no significant upregulation of canonical TGF-β/Smad target genes such as *Serpine1, Smad7*, or *Id3* in mPC61 treated cultures compared to isotype control. Gene set enrichment analysis using the Hallmark “TGF-β signaling” pathway likewise showed no enrichment in the mPC61-treated samples. Collectively, these results demonstrate that while activation of the MACS pathway via mPC61 enhances iTreg differentiation, this effect occurs without evident PC61-induced increased TGF-β/Smad2/3 signaling.

**Figure 6.**
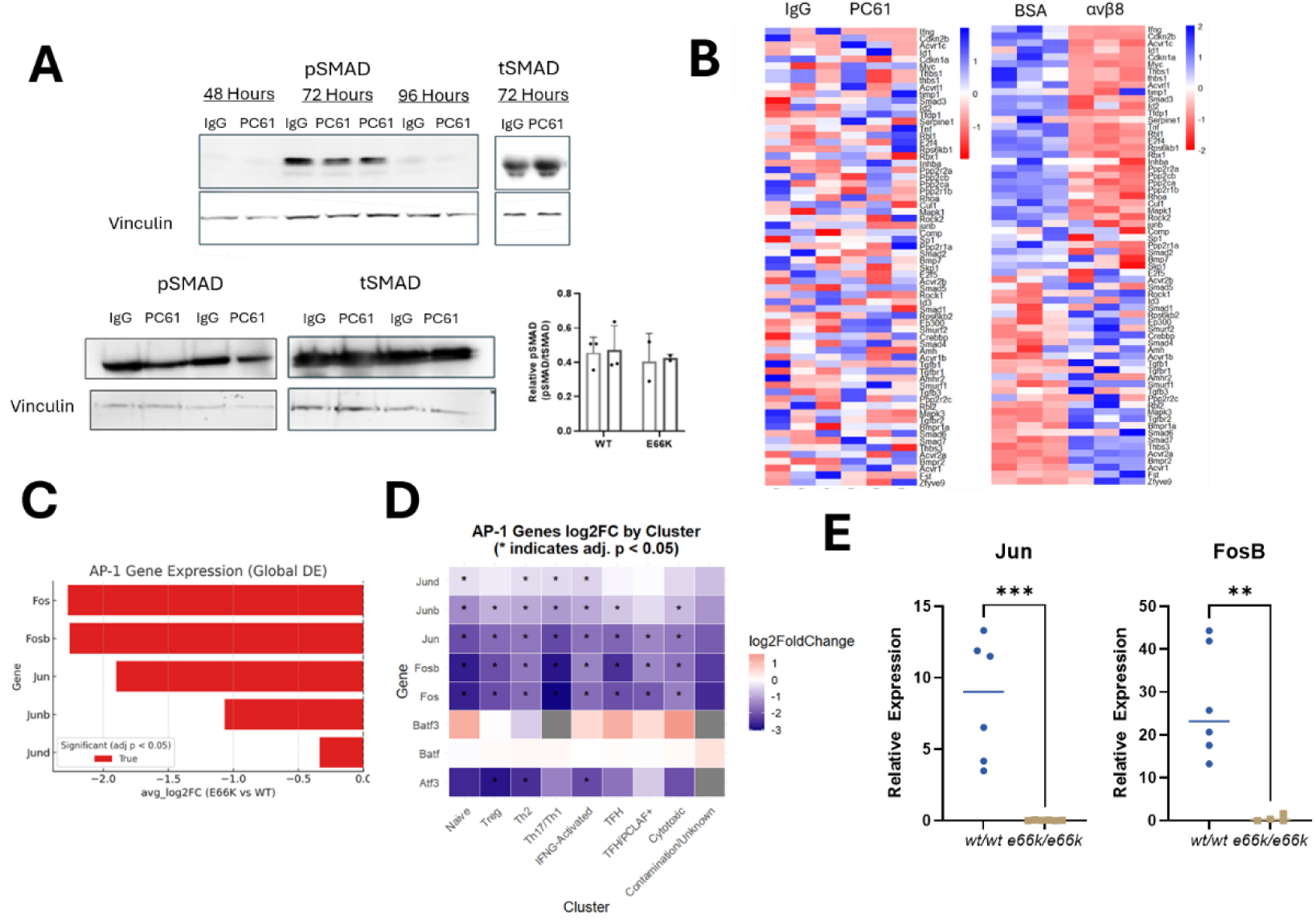
Reduced AP-1 family gene expression in *Il2^E66K/E66K^* T cells. **(A)** Immunoblot analysis of phospho-Smad2/3 and total Smad following iTreg induction in the presence of mPC61 or Isotype control (n = 3) and of WT vs *Il2*^E66K/E66K^ cells following iTreg induction under the same conditions (n = 2) Bars indicate mean ± SE.; dots represent independent experiments. **(B)** Heat maps of TGF-β–responsive gene expression programs in WT iTregs determined by bulk RNA sequencing (n = 3). **(C)** Average log_2_ fold change of AP-1 family gene expression in *Il2*^E66K/E66K^ versus WT cells across the full CD4^+^ single-cell dataset from >24week old mouse mesenteric lymph nodes. **(D)** Heat map showing pseudobulk log_2_ fold changes for AP-1 family members across indicated CD4⁺ T-cell clusters; differential expression significance was assessed using DESeq2. Asterisks denote adjusted P < 0.05. (E) Quantitative PCR analysis of AP-1 gene expression in CD4⁺ T cells isolated from mesenteric lymph nodes (n = 3) of >24 week old mice of the indicated genotype; significance was assessed by two-tailed t-test (n = 3).

### Global reduction of AP-1 transcription factors suggests a mechanism for defective pTreg differentiation in *Il2^E66K/E66K^* mice

Global differential expression analysis of the single-cell transcriptomic data from mesenteric lymph node CD4^+^ cells revealed a striking, global reduction in AP-1 family transcription factors (Fos, Fosb, Jun, Junb) (Figure 6C) in *Il2^E66K/E66K^* mice. Per cluster pseudobulk analysis revealed that these differences were present across nearly all CD4⁺ T cell clusters in *Il2^E66K/E66K^* mice (Figure 6D). This observation was independently confirmed by qPCR analysis of CD4⁺ T cells isolated from mesenteric lymph nodes (Figure 6E). In addition to pSmads, AP-1 components bind the conserved non-coding sequence 1 (CNS1) enhancer of the Foxp3 locus(*29*) and the effect of MACS on AP1 transcription factors could therefore account for MACS effect on Treg differentiation and function.

## DISCUSSION

The capacity of a human IL-2 dependent CD25-chemokine receptor complex to drive *in vitro* Treg integrin activation defined MACS, an alternative IL-2 signaling pathway. Here we extend these *in vitro* principles to mouse IL-2 MACS and use this information to demonstrate a role for IL-2 MACS *in vivo.* Mice homozygous for a separation-of-function *Il2*^E66K^ mutation that selectively disrupts CD25– CCR7 complex assembly while preserving canonical IL-2–STAT5 signaling exhibited a reduction in the aging-associated increase in Treg abundance in the gut-associated lymphoid tissue (GALT). Single cell RNA-seq analysis of mesenteric lymph nodes from these mice revealed reduced expression of the suppressive cytokine, IL-10 mRNA associated with reduced IL-10 abundance in Tregs and corresponding decreased suppressive function. Furthermore, enforced IL-2 MACS increased *in vitro* Treg differentiation that was impaired in *Il2^E66K/E66K^* CD4^+^ cells. Enforcing or inhibiting MACS modulated the expression of AP1 transcription factors that are known to facilitate transcriptional activation of the Foxp3 locus, but did not influence TGF-β signaling to SMAD 2,3 phosphorylation during iTreg induction. Thus, we show that MACS is critical for sustaining suppressive effector Tregs in GALT with age thereby establishing a physiological role for MACS IL-2 signaling pathway in shaping Treg differentiation and function.

The selective reduction of regulatory T cells in gut-associated lymphoid tissues of aged *Il2*^E66K/E66K^ mice, including a preferential loss of Nrp1⁻ Tregs, suggests that MACS disruption does not compromise early immune development but instead disproportionately affects peripherally-derived regulatory populations. This phenotype closely parallels prior studies in which extrathymic Treg differentiation is selectively impaired while thymic Treg development remains intact(*25, 26*). Deletion of the Foxp3 CNS1 enhancer— a regulatory element required for peripheral induction of Foxp3— has been shown to result in marked reductions in mucosal pTregs while sparing thymic Treg populations and avoiding systemic autoimmunity. The age dependence of the phenotype further argues against an obligatory role for MACS in initial establishment of the Treg lineage and is consistent with a progressive defect in peripheral maintenance and/or functional maturation. Several age-associated processes could amplify signaling perturbations: peripheral Treg induction has been reported to decline with age(*30*), and age-related immune remodeling can alter the balance of signals that sustain regulatory programs(*31*).

Single-cell transcriptomic analyses and flow cytometry identified an effector Treg subset with reduced expression of *Il10* and other suppressive markers in *Il2*^E66K/E66K^ mice, and this defect was mirrored functionally in co-culture suppression assays in which *Il2*^E66K/E66K^ Tregs were significantly less effective at inhibiting responder CD4⁺ T-cell proliferation than wild-type Tregs. This phenotype is conceptually aligned with our previous work demonstrating that activation of the MACS pathway using the antibody PC61 enhances Treg suppressive function and increases IL-10 expression(*9*), suggesting that genetic disruption of MACS produces the reciprocal transcriptional and functional outcomes. Reduced IL-10 expression within this subset is likely to be biologically consequential, as IL-10 is a central mediator of immune regulation in mucosal tissues and the central nervous system(*32*), and has been repeatedly implicated in limiting inflammatory pathology in these environments(*33*). Consistently, conditional knockout of IL-10 within Tregs results in tissue-specific inflammation at barrier sites such as the gut, without observed systemic autoimmunity(*33*).

*In vitro*, PC61 enhanced iTreg differentiation in wild-type cultures, whereas this effect was diminished in *Il2*^E66K/E66K^ cells, consistent with a requirement for MACS functionality in this setting. The ability of PC61 to enhance Foxp3 induction *in vitro* has been reported previously; Mannie and colleagues hypothesized that anti-CD25 antibodies can bias mixed T cell cultures, thereby favoring CD25^high^ cells rather than directly promoting regulatory differentiation(*34*). While this mechanism could play a role in our studies, the effect was lost in *Il2*^E66K/E66K^ cultures, consistent with a requirement for MACS-dependent receptor complex formation in addition to CD25 blockade. Additionally, adoptive transfer into Rag1⁻/⁻ hosts revealed a reduced capacity of *Il2*^E66K/E66K^ cells to generate Foxp3⁺ Tregs in vivo, particularly within gut-associated lymphoid tissues. These findings argue that in addition to a simple selection-based mechanism, MACS signaling directly promotes peripheral Treg differentiation in physiological settings.

The PC61-induced iTreg differentiation enhancement occurred without detectable increases in canonical TGF-β/Smad2/3 signaling, suggesting that PC61 does not act by further amplifying Smad-dependent TGF-β activity and may instead engage alternative signaling pathways to facilitate peripheral Treg differentiation programs. One such candidate comprises the AP-1 transcription factors, which are known to bind regulatory elements within the Foxp3 locus, including CNS1, to regulate pTreg differentiation and function (*28, 35–37*) Notably, multiple AP-1 family members were downregulated in *Il2*^E66K/E66K^ cells, raising the possibility that this transcriptional network contributes to downstream regulatory programs influenced by MACS. Briefly describe our findings regarding AP-1

The signals that govern peripheral Treg differentiation *in vivo* remain incompletely defined. Our iTreg experiments were performed under conditions designed to model tissue-relevant induction, using αvβ8-coated plates rather than exogenous TGF-β, reflecting prior work demonstrating that αvβ8 expressed by intestinal dendritic cells and stromal populations activate latent TGF-β and promotes extrathymic Foxp3 induction in the gut(*38–40*). More broadly, αvβ8-dependent pathways have emerged as key regulators of tissue Treg programs, and recent work in fatty liver has shown that the AP-1 family transcription factor ATF4 drives transcriptional programs that promote αvβ8-mediated TGF-β activation and are required for the differentiation and accumulation of effector Tregs in non-lymphoid tissues(*41*). Together, these observations place MACS within a landscape of tissue-specific signaling pathways that shape peripheral regulatory immunity.

Limitations: Although the IL-2(E66K) mutation was designed to preserve IL-2 binding to the trimeric canonical IL-2 receptor and did not impair signaling through the JAK–STAT5 pathway, there remains a theoretical possibility that it affects unknown IL-2 functions beyond blocking MACS. In addition, because CCR5 is up-regulated on activated Tregs and IL-2(E66K) does not prevent CCR5-mediated MACS(10), the *Il2^E66K/E66K^*mouse could not be used to elucidate the effects of MACS loss in models of inflammation.

## MATERIALS AND METHODS

### Study design

The aim of this study was to determine the physiological role of noncanonical interleukin-2 (IL-2) signaling in regulatory T cell differentiation and function. Il2^E66K/E66K^ mice, in which IL-2–dependent complex formation is disrupted while canonical IL-2–STAT5 signaling is preserved, were compared with wild-type littermate controls. To achieve this aim, the following experiments were performed; regulatory T cell abundance, phenotype, transcriptional state, differentiation, and suppressive function were evaluated using flow cytometry, as well as in vitro differentiation and suppression assays, adoptive transfer experiments, immunoblotting, quantitative PCR, bulk RNA sequencing, and single-cell RNA sequencing. Experimental units were individual mice or independently performed cell-culture experiments, as indicated in the figure legends. Sample sizes and numbers of biological and technical replicates are reported in the corresponding figure legends.

### Generation of Il2^E66K/E66K^ strain

A ribonucleoprotein (RNP) complex consisting of 4 µM Alt-R CRISPR RNA (5’-CTGAAGATCTTTCAATTCTG-3’), 4 µM tracrRNA, and 3 µM HiFi Cas9 nuclease V3 (Integrated DNA Technologies, IDT; Coralville, IA) along with a 4 µM single-stranded oligodeoxynucleotide (ssODN) donor template (5’-AGCTTATTACTCCTCTAGGCCACGAAATTAAAGGACCTCCAATGCCTAGAA-3’) was electroporated into C57BL/6J (B6J) zygotes (Jackson Laboratory, stock #000664; Bar Harbor, ME). Surviving embryos were implanted into pseudopregnant ICR females, and pregnancies were carried to full term. Tissue biopsies for genomic DNA extraction were collected from G0 founder pups between postnatal days 7 and 10 to determine the *Il2^WT/E66K^* founders. Mice were genotyped by PCR using forward primers 5’-GTGGCCTAACAGTCAAGGTG-3’ and reverse primers 5’-GACCAGAAATGGCAAGCCAC -3’ followed by Sanger sequencing using forward primers 5’-GTGGCCTAACAGTCAAGGTG -3’ for E66K mutation.

Founder *Il2^WT/E66K^* mice were propagated on a C57BL/6 background and backcrossed to Foxp3^GFP^ strain to obtain *Il2^E66K/E66K^Foxp3^GFP^*. Experimental cohorts were generated from Il2^WT/E66K^ intercrosses, yielding Il2^WT/WT^, Il2^WT/E66K^, and Il2^E66K/E66K^ littermates. Sex-matched *Il2^wt/wt^* littermates were used as controls for *Il2^E66K/E66K^* mutant animals in all experiments. Mice were housed under specific pathogen– free conditions at the University of California San Diego. All animal procedures were approved by the University of California San Diego Institutional Animal Care and Use Committee (IACUC) and were performed in accordance with institutional guidelines and the National Institutes of Health Guide for the Care and Use of Laboratory Animals.

Potential CRISPR/Cas9 off-target sites for the Il2^E66K^ guide RNA were predicted *in silico* using the IDT CRISPR-Cas9 guide RNA design checker. Candidate off-target loci were prioritized based on predicted off-target score, mismatch number, and genomic location. The 10 highest scoring predicted off-target sites were PCR-amplified from genomic DNA and assessed by Sanger sequencing. No protein-altering variants were identified.

### Immunophenotyping

Spleen, peripheral lymph nodes, mesenteric lymph nodes, Peyer’s patches, and colon from adult (8-12 weeks old) and Aged (24+ weeks old) mice from *Il2^E66K/E66K^*and *Il2^wt/wt^* littermate controls were collected for immunophenotyping. Isolated cells were stained with 1 ug/mL of antibodies for 30 minutes at room temperature in the dark and measured on Accuri C6+ flow cytometer (BD Biosciences). Antibodies used were: FITC-CD4 (Biolegend #100510), PerCP-CD4 (Biolegend #116012), PE-CD8 (Biolegend #100708), PE-CD25 (Biolegend #102008), PE-CCR7 (Biolegend #120106), APC-Nrp1 (Biolegend #145206).

### Treg suppression assay

CD4⁺ CD62L^hi^FOXP3-GFP− responder cells were FACS-sorted and labeled with 5 µM of eFluor670 cell proliferation dye (Invitrogen) per manufacturer’s protocol. In a round-bottom 96-well plate, 5 × 104 stained responder cells/well were added to wells each containing 5 µg/mL anti-CD3 (BioLegend, #100301), and 3 × 10^3^ bone-marrow derived dendritic cells (BMDCs) that had been stimulated for 7 days with Granulocyte Macrophage Colony-stimulating Factor (GM-CSF, BioLegend #576302) in the presence or absence of 6.25 x 10^3^ FACS-sorted Tregs (CD4⁺ FOXP3-GFP^+^)/well. The plates were incubated at 37°C and 5% carbon dioxide (CO2) for 4 days and proliferation of responder cells was analyzed by dilution of eFluor670 fluorescence with a BD Accuri C6+ flow cytometer. The Division index of responder cells was calculated using FlowJo with eFluor670 MFI, and % suppression was calculated as:

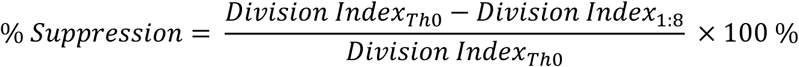

Where Th0= Division Index in the absence of Tregs and 1:8=Division Index in the presence of 6.25 x 103 Tregs/well. Triplicate determinations were obtained in each experiment and the results shown are calculated from four experiments so performed.

### Quantifying supernatant Il-2 via enzyme-linked immunosorbent Assay (ELISA)

Total CD4⁺ T lymphocytes from 8-12 week-old *Il2^E66K/E66K^Foxp3^GFP^* and *Il2^wt/wt^Foxp3^GFP^* littermate controls were isolated from spleens using Mojosort CD4 T cell isolation kit (BioLegend #480033) per Manufacture’s protocol. 1 x 10^5^ CD4⁺ T cells were cultured in a 96-well plate coated with 1ug/ml of anti-CD3 (BioLegend, Cat# 100340) and Anti-CD28 (BioLegend, Cat# 102116), in complete media RPMI 1640 (Gibco 11875093supplemented with heat-inactivated Fetal Bovine Serum (FBS, Gibco, # 16140071), pen/strep Gibco, #15140122 ) non-essential amino acids (Gibco, #11140), sodium pyruvate (Gibco #11360020), and HEPES (Gibco #15630080). After 3 days, supernatants were collected for IL-2 ELISA (Biolegend, #431004). ELISA was performed per manufacturer’s instructions and measured using Mini ELISA Plate Reader (Biolegend, #423555) at 450 nm. Concentrations of IL-2 from supernatants were calculated according to the luminescence intensity from the standard samples prepared in parallel.

### Intracellular staining of Il-2

Total CD4⁺ T cells from 8-12 week-old from *Il2^E66K/E66K^* and *Il2^wt/wt^* littermate controls were isolated from spleens using Mojosort CD4 T cell isolation kit (BioLegend #480033) per Manufacture’s protocol. 1 x 10^5^ CD4⁺ T cells were cultured in a 96-well plate in complete media, RPMI 1640 (Gibco 11875093) supplemented with heat-inactivated Fetal Bovine Serum (FBS, Gibco, #16140071), pen/strep Gibco, #15140122) non-essential amino acids (Gibco #11140), sodium pyruvate (Gibco #11360020), and HEPES (Gibco #15630080). After 3 days, cells were washed with 1X PBS (phosphate buffered saline) followed by incubation with Cell Activation Cocktail with Brefeldin A (Biolegend #423303) for 4 hours in 37°C. After the incubation, cells were stained for live/dead (ZombieGreen, 1:1000, Biolegend # 423111) and APC-CD4 (Biolegend # 100411). Then, cells were fixed and permeabilized with Foxp3/Transcription Factor Staining Buffer Set (eBioscience, 00-5523-00) per manufacturer’s instructions. Cells were stained with PE-mouse IL-2 (Biolegend #503807) and analyzed on Accuri C6+ (BD Biosciences).

### STAT5 Phosphorylation

Foxp3-GFP^+^ Tregs were plated in each well of a 96-well plate and incubated with indicated concentrations of IL-2 in PBS. Cells were stimulated for 30 min at 37°C and immediately fixed and permeabilized using the Transcription Factor Phospho Buffer Set (BD Biosciences #563239). Cells were then washed and incubated with APC-anti STAT5 (1:50, BD Biosciences #612599) for 1 h on ice in the wash buffer. Cells were then washed twice, once with the wash buffer and next with PBS, and MFI was determined on Accuri C6+ flow cytometer (BD Biosciences). Triplicate measurements were made in each experiment, and each experiment was independently replicated at least three times with similar results.

### Soluble Ligand Binding Assay

Cells were plated in a 96-well plate and incubated with 5 μg/mL MAdCAM-1 Fc (R&D Systems #6056-MC) for 30 min at 37°C in HBSS with Ca/Mg (Gibco #14-025-092) with the following conditions: 0.5 μg/mL IL-2, 10 μg/mL IgG control or PC61. In parallel 10 mM EDTA (Invitrogen #AM9260G) was added in each condition to measure non-specific binding. Cells were then immediately washed with 1X HBSS with Ca/Mg, followed by incubating with APC-anti-human IgG Fc (1:25, BioLegend #410714) for 1 h on ice. Cells were washed in 1X HBSS with Ca/Mg, and geometric mean fluorescence intensity (MFI) was determined on an Accuri C6+ flow cytometer (BD Biosciences). Geometric mean fluorescence intensity (MFI) of EDTA groups were subtracted from MFI without EDTA groups to report ligand binding (MFI EDTA). Triplicate measurements were made in each experiment, and each experiment was independently replicated at least three times with similar results.

### Differentiation of Tregs in vitro

Splenocytes were isolated *Il2^wt/wt^Foxp3*^GFP^ *or Il2^E66K/E66K^Foxp3*^GFP^ male mice by mechanical dissociation using a sterile tissue homogenizer, followed by red blood cell lysis with ACK lysis buffer (Gibco, Cat# A10492). Cells were filtered through a 100 µm cell strainer and CD4 naïve cells purified by magnetic separation using the MojoSort™ Mouse CD4 Naïve T Cell Isolation Kit (BioLegend, Cat# 480040) and a biotin-conjugated anti-CD25 antibody (PC61; BioLegend, Cat# 102004) to deplete Tregs. Purified naïve CD4⁺ T cells were seeded at 1 × 10^5^ cells per well in a 96-well plate pre-coated with anti-CD3 (2µg/mL; BioLegend, Cat# 100340), anti-CD28 (2µg/mL; BioLegend, Cat# 102116), and recombinant αvβ8 protein (1 µg/mL; R&D, Cat#4135-AV-050). Cells were cultured in complete RPMI 1640 Glutamax medium (Gibco, Cat# 61870036; with 25mM HEPES (Gibco, Cat# 15630080), 1% Sodium Pyruvate (Gibco, Cat# 11360020), 1% NEAA (Gibco, Cat# 11140) and 0.1% bME (Gibco, Cat# 21985023). Cultures were supplemented with 6 ng/mL murine WT or E66K IL-2 for 4 days at 37°C with 5% CO₂ and treated with 5ug/ml mPC61 or IgG1 (BioXCell, #BE0083).

### Single Cell RNA Sequencing

Single-cell RNA sequencing (scRNA-seq) was performed with assistance from the UC San Diego IGM Genomics Center. CD4⁺ T cells were isolated from mesenteric lymph nodes by magnetic-activated cell sorting (MACS) as described above. Cells were assessed for viability and abundance prior to library preparation. Single-cell libraries were generated using the 10x Genomics Chromium(*42*) platform following the manufacturer’s protocols. Single-cell suspensions were loaded onto Chromium microfluidic chips to generate gel bead–in-emulsions (GEMs), followed by reverse transcription, cDNA amplification, and library construction. Libraries were sequenced on an Illumina Novaseq at the UCSD core facility.

Raw sequencing data were processed using Cell Ranger (10x Genomics) for demultiplexing, alignment to the mouse reference genome, and generation of gene–barcode count matrices. Downstream analyses were performed in R using Seurat(*43*). Cells were filtered based on standard quality-control criteria, including the number of detected genes, UMI counts, and the proportion of mitochondrial transcripts. Data were normalized, highly variable genes were identified, and dimensionality reduction was performed by principal-component analysis followed by Uniform Manifold Approximation and Projection (UMAP)(*44*). Graph-based clustering was used to define transcriptionally distinct populations, which were annotated using established marker genes. For differential gene-expression analysis, pseudobulk count matrices were generated by aggregating reads from cells within each cluster for individual biological replicates. Differential expression testing between genotypes was performed using DESeq2 (*45*), with significance determined after correction for multiple comparison testing.

Across three biological replicates per genotype, wild-type samples yielded an estimated 136,100 cells with 94.7% of reads mapping to barcodes, a mean of 1,810 reads per cell, and a median of 526 genes detected per cell. *Il2*^E66K/E66K^ samples yielded an estimated 103,935 cells with 94.2% of reads mapping to cellular barcodes, a mean of 2,153 reads per cell, and a median of 615 genes detected per cell. Total genes detected across datasets were 22,681 and 22,264, respectively, indicating comparable sequencing depth and library complexity between groups.

### RNA Extraction

Total RNA was isolated from cell pellets using the Qiagen RNeasy Micro Kit (CAT#74004) according to the manufacturer’s instructions, including on-column DNase I treatment to remove genomic DNA contamination. Briefly, cell pellets were lysed in the supplied buffer, and RNA was purified using silica-membrane spin columns with sequential washes and elution in RNase-free water. Purified RNA was stored at −80 °C until downstream analysis.

### Quantitative PCR

RNA was isolated as described above and quantified prior to analysis. One-step reverse transcription quantitative PCR (RT–qPCR) was performed using the ZymoScript One-Step RT-qPCR Kit (R3014; Zymo Research) according to the manufacturer’s instructions. Reactions were assembled using gene-specific primers and run using the manufacturer-recommended cycling conditions. Primer sequences were as follows: Jun forward 5′-GCTGGAAGAAGGGCTGTTG-3′ and reverse 5′-CAGATTCAAAGTTGGAAGGAGAC-3′; Fosb forward 5′-ACCCTCTGCCGAGTCTCAATAT-3′ and reverse 5′-GCCACTGCTGTAGCCACTCAT-3′. All samples were run in technical triplicate. Ct values were calculated using instrument software, and relative expression was determined using the ΔΔCt method after normalization to glyceraldhyde phosphate dehydrogenase (GAPDH). No-template controls were included for each primer pair to assess contamination

### Western Blotting

Cells were prepared and stimulated as described above for an iTreg assay. At the indicated time points, cells were lysed directly in SDS sample buffer containing protease and phosphatase inhibitors (Rocher, Cat# 589297001), centrifuged to remove insoluble material, and briefly sonicated (30 s) to reduce viscosity. Clarified lysates were heated for 5 min and resolved by SDS–PAGE on 4–12% gradient polyacrylamide gels (Life Tech; XP04200BOX). Proteins were transferred to nitrocellulose membrane by wet transfer in Tris–glycine buffer containing ethanol, and transfer efficiency was verified by Ponceau staining. Membranes were blocked in 5% BSA in TBST for at least 30 min and incubated overnight at 4 °C with primary antibodies diluted 1:1,000 in 5% BSA/TBST, including phospho-Smad2/3 (CAT# ab272332) and total-Smad2/3 antibodies. After washing, membranes were incubated with HRP-conjugated secondary antibodies (ThermoFisher, Cat#31460) for 1 h at room temperature. Blots were developed using enhanced chemiluminescence substrate (ThermoFisher, Cat#A38554) and imaged on a GelDoc system.

### Flow Cytometry

#### GFP-based quantification of Tregs

For quantification of Foxp3⁺ Tregs, single-cell suspensions were analyzed based on Foxp3-driven GFP expression. Cells were stained with propidium iodide (PI) to exclude dead cells and acquired on a BD Accuri flow cytometer. Data were analyzed by sequential gating on singlets and PI⁻ live cells, followed by quantification of GFP⁺ events.

#### Multicolor flow cytometry

For multicolor immunophenotyping, samples were acquired at the La Jolla Institute for Immunology (LJI) Flow Cytometry Core Facility. Live/dead discrimination was performed using Ghost Dye Red 710 (Cell Signalling, CAT# 90232). Surface staining was performed using antibodies against CD73 (PerCP-Cy5.5; TY11.8/CAT#127220 ), NRP1 (PE; CAT#145206), CD69 (PE-CF594; H1.2F3/CAT# 562455), LAG-3 (APC-Fire 750; C9B7W/CAT# 125231), TIGIT (BV421; IG9/CAT# 565270), and CD25 (BUV737; PC61/CAT# 569602). For intracellular staining, cells were fixed in 2% paraformaldehyde and permeabilized using a commercial Foxp3/transcription factor staining kit (Invitrogen; CAT# 00-5523-00) prior to staining for IL-10 (APC; JES5-16ER/CAT# 554468). Foxp3 expression was tracked via GFP. Acquired samples were analyzed on FlowJo v10.

#### Co-Immunprecipitation

For heparin sulfate stimulation, Membrane fractions from HEK293T cells expressing HA-mCD25 were prepared by differential centrifugation as described previously and stimulated at 37 °C for 30 min with heparan sulfate (HS, 5 μg/mL) together with IL-2 (WT or E66K, 1 μg/mL). For PC61-mediated engagement, the cells were directly stimulated by PC61 (5 μg/ml) and mouse IL 2 (1 μg/ml). Cells were lysed on ice for 15 min in lysis buffer (50 mM Tris Cl, pH 7.4; 150 mM NaCl; 0.5% NP 40; 0.5 mM CaCl₂; 0.5 mM MgCl₂; protease inhibitor cocktail, mini tablet; MedCamExpress, HY K0011). Soluble lysates were clarified by centrifugation at 12,000 rpm for 15 min. Supernatants were incubated overnight at 4 °C with anti HA monoclonal antibody (12CA5) bound to protein G beads (Genscript, L00209). After three washes with lysis buffer, bead associated proteins were eluted and analyzed by SDS PAGE. Coprecipitated CD25 and CCR7 were detected by immunoblotting using an anti HA antibody (BioLegend, 901513) and an anti-CCR7 antibody (Novus Biologicals, NBP2-67324). Band intensities were quantified using an Odyssey CLx infrared imaging system (LI COR Biosciences).

#### Adoptive transfer experiment

Splenocytes were isolated from *Il2*^wt/wt^ *Foxp3-GFP* and *Il2*^E66K/E66K^Foxp3GFP mice by mechanical dissociation and ACK lysis as described. Cells from *Foxp3-GFP* mice were used as a source of naïve CD4⁺ T cells, Foxp3-GFP⁻ mice were used as a source of Tregs. CD4⁺ T cells were enriched by magnetic separation (MojoSort) and subsequently purified by FACS sorting. Cells were stained as described above, resuspended in complete media, and naïve CD4⁺ T cells and Tregs were combined at a 10:1 ratio. Cells were injected retro-orbitally into recipient *Rag1⁻/⁻* mice, which were harvested 60 days post-transfer for analysis.

#### Statistical Analysis

Statistical analysis was performed using PRISM software (version 9.00, GraphPad Software), and all datasets were checked for Gaussian normality distribution. Data analysis was performed using two-tailed t-test, one-way ANOVA or two-way ANOVA followed by Tukey’s correction as indicated in the Figure legends. The resulting P values are indicated as follows: NS: not significant, *: p <0.05; **: p<0.01. Plotted data are the mean ± SEM of at least three independent experiments.

### Supplementary Materials

**Fig. S1.** Peripheral blood counts in *Il2^E66K/E66K^* mice and littermate controls.

**Fig. S2.** Single-cell RNA-sequencing quality-control metrics.

**Fig. S3**. Marker-gene expression used to annotate CD4⁺ T cell clusters.

## Author contributions

Conceptualization: AM, SHJK, LFL, CJ, MG, MALR, MHG

Methodology: AM, SHJK, MHG, HSL

Investigation: AM, SHJK, MT, BL, SC, HSL, ZZ, MHFL

Formal analysis: AM, SHJK, HSL, MHG

Visualization: AM, SHJK, HSL, MHG

Resources: LFL, CJ, MG, MHG

Supervision: MHG

Writing – original draft: AM, SHJK, MHG

Writing – review & editing: AM, SHJK, MHFL, MT, BL, SC, HSL, LFL, ZZ, CJ, MG, MALR, MHG

## Data and materials availability

The single-cell RNA-sequencing data generated in this study will be deposited in the Gene Expression Omnibus (GEO) under accession number XXXXX upon publication. Processed count matrices and metadata will be made available through GEO and upon reasonable request from the corresponding author.

