## Supplemental Figure 1, Supplemental Figure 2, Supplemental Figure 3 for "Non-canonical IL-2 signaling facilitates peripheral regulatory T cell differentiation"

### Supplementary 1

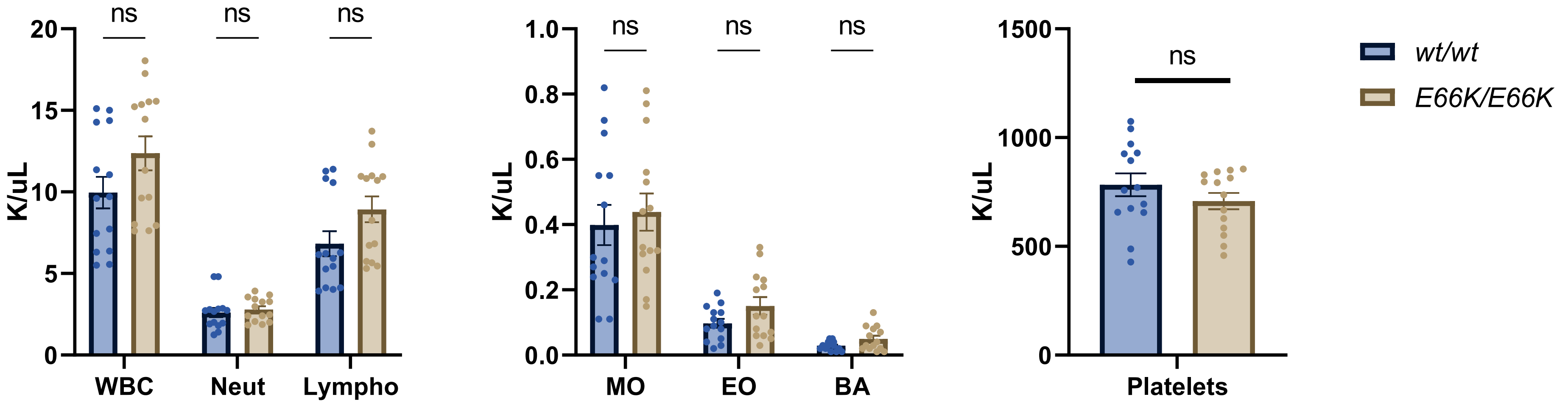

### Supplementary 2

| Sample | Genotype | Cells retained | Median genes per cell | Median UMIs per cell |
|---|---|---|---|---|
| **WT-1** | WT | 54148 | 436 | 568 |
| **WT-2** | WT | 35129 | 633 | 931 |
| **WT-3** | WT | 45098 | 508 | 677 |
| **K-1** | E66K | 43241 | 469 | 592 |
| **K-2** | E66K | 25104 | 729 | 1026 |
| **K-3** | E66K | 33622 | 656 | 910 |

### Supplementary 3

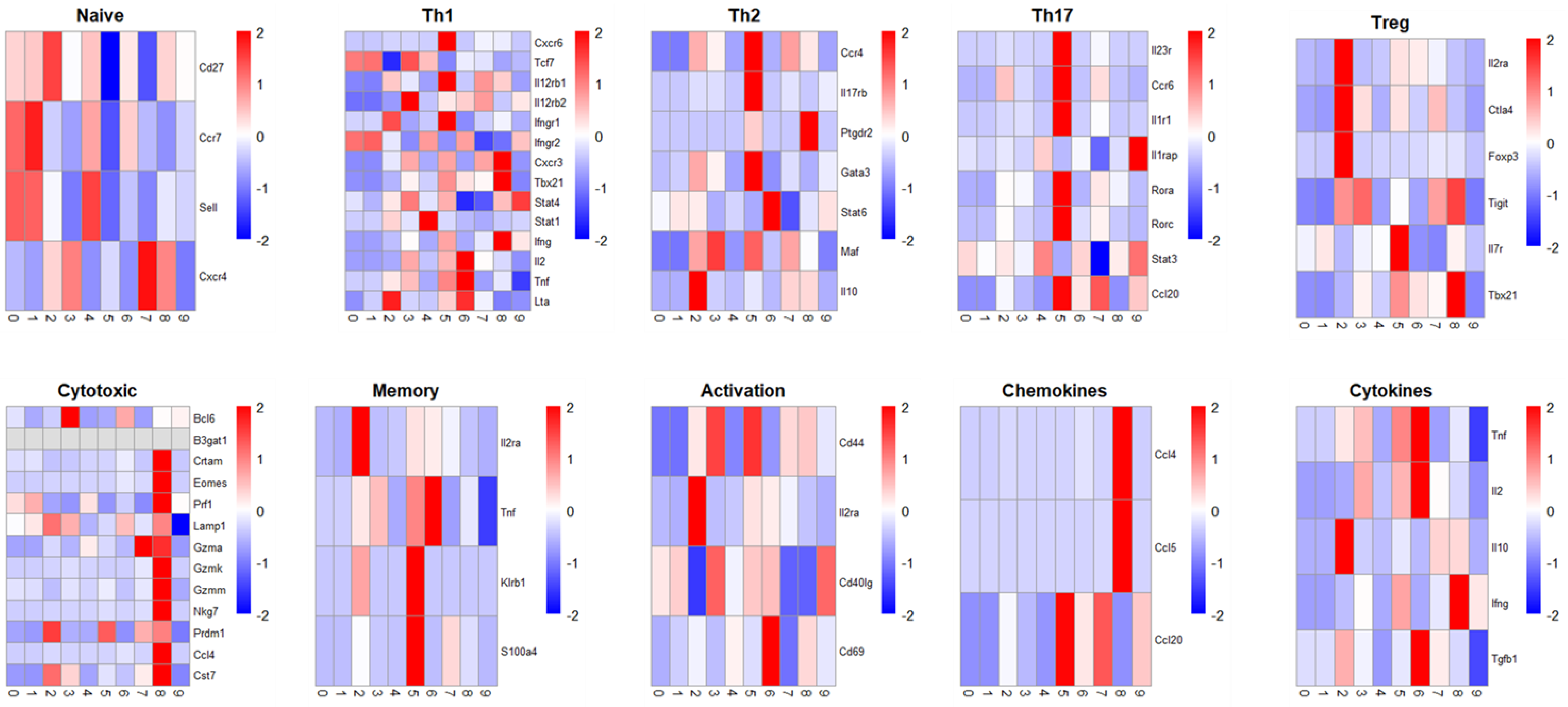
